# Bearded or smooth? Awns improve yield when wheat experiences heat stress during grain fill

**DOI:** 10.1101/2023.02.27.530138

**Authors:** Noah DeWitt, Jeanette Lyerly, Mohammed Guedira, James B. Holland, Brian P. Ward, J. Paul Murphy, Richard E. Boyles, Mohamed Mergoum, Md. Ali Babar, Ehsan Shakiba, Russel Sutton, Amir Ibrahim, Vijay Tiwari, Nicholas Santantonio, David A. Van Sanford, Kimberly Howell, Jared H. Smith, Stephen A. Harrison, Gina Brown-Guedira

## Abstract

The presence or absence of awns – whether wheat heads are ”bearded” or ”smooth”– is the most visible phenotype distinguishing wheat cultivars. Previous studies suggest that awns may improve yields in heat or water-stressed environments, but the exact contribution of awns to yield differences remains unclear. Here we leverage historical phenotypic, genotypic, and climate data to estimate the yield effects of awns under different environmental conditions over a 12-year period in the Southeast US. Lines were classified as awned or awnless based on sequence data, and observed heading dates were used to associate grain fill periods of each line in each environment with climatic data and grain yield. In most environments, awn suppression was associated with higher yields, but awns were associated with better performance in heat-stressed environments more common at southern locations. Wheat breeders in environments where awns are only beneficial in some years may consider selection for awned lines to reduce year-to-year yield variability, and with an eye towards future climates.

## Introduction

Humans rely more on wheat (*Triticum aestivum L*.) than any other crop species for both direct calories and protein (FAO, 2020). Despite its global range, wheat is more sensitive to heat and drought stress than other grains like maize (*Zea mays* L.) and sorghum (*Sorghum bicolor* L.), in part due to a lower water use efficiency compared to its C4 relatives (Aggarwal and Sinha, 1983). Heat stress hampers plants’ ability to fill grain, through decreasing sucrose levels and the overall duration of grain fill, and through deactivating the wheat starch synthase enzyme responsible for assimilating carbohydrates into developing grain (Bhullar and Jenner, 1985; Hawker and Jenner, 1993; Guedira and Paulsen, 2002). The simplest way to breed for drought and heat tolerance in areas where these stresses are common is to plant yield trials in those environments. However, as climate change threatens higher temperatures and altered precipitation patterns in areas without those stresses, an understanding of the genetic and physiological basis of drought and heat stress response will help breed for future conditions.

A principal morphological trait implicated in wheat drought and heat stress response is the presence or absence of awns or ”beards”. Awns are thin, rigid extensions of the lemma that give wheat heads a brush-like appearance. In US germplasm, the presence or absence of awns is determined by the *B1* (*Tipped 1*) awn suppressor gene near the end of the long arm of chromosome 5A (Kato et al., 1998; DeWitt et al., 2020). Awn suppression results from over-expression of *B-A1* which encodes a C2H2 zinc finger (Huang et al., 2020; DeWitt et al., 2020). Suppression of awns by *B1* produces awn lengths ranging from a totally awnless phenotype to an ”awnletted” phenotype where most spikelets lack awns, but short awns are present near the tip of the spike (Watkins and Ellerton, 1940). Outside of US germplasm, additional genes *B2* (*Tipped 2*) and *Hd* (*Hooded*) may produce a semi-awnless phenotype alone or a completely awnless phenotype in combination (Watkins and Ellerton, 1940). Within awnletted *B1* US wheat, there remains quantitative variation for tip awn length. For simplicity we will refer to all classes of awnless and awnletted phenotypes as ”awnless”.

As a typically monogenic and visually striking trait, studies on awns constitute some of the earliest in genetics research. In Ohio field trials, Hickman (1889) found that ”bearded”cultivars out-yielded ”smooth”cultivars 40.5 to 37.5 bu acre^−1^ (2.72 to 2.52 Mg ha^−1^). As early as 1937, it was ”generally agreed that in certain regions the awned cultivars out-yield awnletted or awnless ones”, and that ”awned cultivars have been selected and grown in regions of limited rainfall, for awnless types are generally not well-adapted to such conditions”(Gauch, 1937). The distribution of awned and awnless types in wheat lines globally supports this observation (Börner et al., 2005; DeWitt et al., 2020). Anecdotally, breeders targeting environments that are warmer and dryer during the late growing season are disinclined to select awnless lines, while breeders in cooler climates may prefer to select awnless lines.

Research into the yield effects of awns, and the role of heat and drought stress in those effects, has centered on the photosynthetic potential of awn tissue (Grundbacher, 1963; Tambussi et al., 2005; Amjad Ali et al., 2010; Motzo and Giunta, 2002; Li et al., 2006). The movement of carbohydrates from awns into developing grain is facilitated by their close physical proximity, and awns may consist of up to half the photosynthetic area of the late-season wheat canopy. As might be expected for a trait which has an environmentally conditional effect on yield, estimates of the yield effects of awns have shown some cases where awned lines out-perform their awnless counterparts (Vervelde, 1953; Weyhrich et al., 1994; Motzo and Giunta, 2002; Martin et al., 2003), but also some where no significant differences are identified (Foulkes et al., 2007; Rebetzke et al., 2016; Sanchez-Bragado et al., 2020). Awns are more effective at photosynthesizing in water-limited conditions, using less water per quantity of CO_2_ assimilated, and contribute more carbohydrates relative to other tissues in water-limited and heat-stressed conditions (Evans et al., 1972; Blum, 1986; Weyhrich et al., 1995; Li et al., 2006; Maydup et al., 2014). Awn tissue often senesces last, and has a higher temperature optimum for photosynthesis than flag leaf and spike tissue, which may contribute to grain yield in heat-stressed conditions (Blum, 1986). At the same time, the adaptive advantage of awns in heat-stressed environments may relate directly to the ability of awns to help cool ears by transferring heat into the atmosphere. In studies of canopy temperature, awns cool the head during grain fill (Ferguson, 1975; Motzo and Giunta, 2002; Rebetzke et al., 2016), which may reduce stress in a crop sensitive to heat during this growth stage. The ability of awns to cool canopy temperatures and more efficiently photosynthesize in water-limited conditions may explain the perceived yield benefits of awned lines in warmer and drier climates. In general, awns tend to be associated with an increase in grain weight at the expense of grain number (Rebetzke et al., 2016; DeWitt et al., 2020; Sanchez-Bragado et al., 2020). The increase in total seed number associated with *B1* awn suppression may explain the prevalence of awnless varieties in some environments, as increased seed number may lead to greater yields in environments where plants are not source-limited or commonly stressed during grain fill. *B1* increases seed number through either increasing the number of spikelets per spike or grains per spikelet (Dewitt et al., 2022), effects which may be caused by either resource allocation towards awns in the developing spike or to other developmental effects of an increase in expression of the *B1* transcription factor beyond awn suppression.

Previous studies on the effect of awns on grain yield have focused on comparisons of near-isogenic lines or a small number of lines in a limited sample of environments. To this point no studies have directly estimated reaction norms for the interaction of awn status with environmental conditions. Here we use a large historic data set consisting of real breeding lines where the frequency of *B1* is close to 50% to estimate the allele by environment effect of *B1*. Previous genotyping of these lines and identification of a SNP near *B-A1* predictive of awn status allows us to both classify historic lines into awned and awnless, and to control for population structure with genotypic data when estimating allele effects. Observed heading date was used to estimate grain fill periods for each line in each environment, and weather data collected from public databases was averaged across this period to test the interaction of *B1* with environmental variables. We use the estimated effect of awns on yield in interaction with maximum grain fill temperatures to investigate the utility of selection for or against awned lines in different environments.

## Materials and methods

### Phenotypic data

Data on yield and heading date was collected for plots in cooperative nurseries representing 1,376 late-stage wheat lines submitted by participating public breeding programs in the Southern and Mid-Atlantic winter wheat growing region of the United States. The Southern-most breeding programs consisted of members of the SunGrains^®^ breeding cooperative: Louisiana State University, University of Arkansas, Clemson University, Texas A&M University, University of Georgia, University of Florida, and North Carolina State University. Additional participating programs were located at the University of Kentucky, the University of Maryland, Virginia Polytechnic Institute and State University (Virginia Tech), and the USDA-ARS SEA Plant Science Research unit in Raleigh, NC. Data from three cooperative yield trial nurseries (the Mason-Dixon, Gulf Atlantic Wheat Nursery (GAWN), and SunWheat) were used, to which individual programs annually submitted entries that were planted once at all sites, but typically within only a single year, excluding check lines that were planted across years to estimate year effects (Table 1). The Mason-Dixon generally consisted of entries submitted by the University of Kentucky, University of Maryland, Virginia Tech, USDA-ARS SEA Raleigh, and occasionally the North Carolina State University programs. The GAWN nursery generally consisted of entries from the SunGrains^®^ and Virginia Tech programs, while the SunWheat nursery consisted entirely of SunGrains^®^ lines. Individual trials consisted of two to four replications of each line, and analyzed data consisted of entry-means calculated by collaborators for each line in each site-year combination. Trials were planted in the October or November prior to the harvest year and were at minimum 1.3 m wide and 3.1 m long. Management of plots varied by collaborator depending on state-specific recommendations. Yield was reported in bu acre^−1^ based on whole-plot harvesting and weighing of grain, assuming a consistent test weight of 60 lb bu^−1^ and adjusting for moisture content when appropriate. Heading date values for lines were recorded as the date on which approximately half of wheat heads had emerged from the boot.

**Table 1:** Phenotypic data by nursery. Year ranges, mean number of testing sites and evaluated lines per year, total number of entry means (trial by genotype combinations), and percentage of tested lines with awns are given per nursery.

| Nursery | Years | Mean # Sites | Avg # Lines | Entry Means | Awned |
| --- | --- | --- | --- | --- | --- |
| Mason-Dixon | 2012-2018 | 3.6 | 76.94 | 1,385 | 46% |
| GAWN | 2008-2020 | 5.5 | 58.82 | 4,235 | 52% |
| SunWheat | 2014-2020 | 4.6 | 85.03 | 2,721 | 64% |

### Genotypic data

Tissue samples from nursery lines were submitted to the USDA-ARS Eastern Regional Small Grains Genotyping Lab for genotyping via genome-wide single nucleotide polymorphism (SNP) markers. A genotype by sequencing approach using the PstI and MspI enzymes was performed per Poland et al. (2012) to allow for joint discovery and calling of SNPs. SNP discovery was based on GBS reads from both the nursery lines phenotyped in this study and additional soft red winter wheat (SRWW) lines to capture additional variants that may be at low frequency in the study set. Reads were aligned to version 1.0 of the International Wheat Genome Sequencing Consortium RefSeq assembly of Chinese Spring (https://wheat-urgi.versailles.inra.fr/Seq-Repository/Assemblies) using version 0.7.12 of the Burrows-Wheeler algorithm (Li et al., 2009) via Tassel 5GBSv2 pipeline version 5.2.35 (Glaubitz et al., 2014). The resulting Tassel discovery database was used to call SNPs using reads from the nursery lines, and called SNPs were filtered to exclude markers with more than 50% total missing data, less than 5% minor allele frequency, and greater than 10% heterozygosity given the expected inbreeding of genotyped lines. Beagle version 5.2 was used to impute missing SNPs (Browning and Browning, 2016).

DeWitt et al. (2020) found that in US SRWW germplasm, awn status is a monogenic trait controlled by the *B1* gene. They identified a single GBS marker on chromosome 5A at 698528417 bp (Refseq v1.0 coordinates) predictive of *B1* in 98% of 640 SRWW lines that constitute a subset of the lines in this data set. This SNP was also identified in the SNP discovery database of the full set of SRWW lines, and used to determine allelic status of *B1* in genotypes for which data on awn status was not available.

Covariance between allele frequency of *B1* and population structure could bias estimates of interactions between awns and environmental factors. GBS data was used to estimate realized relationships between nursery lines for later population structure analysis. Markers were first thinned based on LD (*r*^2^ < 0.8), before being coded in terms of the allele dosage of the minor allele. The realized relationship matrix **G** was computed via the Van Raden method (VanRaden, 2008) as 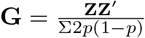, where *p* represents a vector of allele frequencies of all SNP in the (−1, 0, 1) coded marker matrix **M**, and **Z** = **M** -**P**, where **P** is a matrix consisting of the vector *p* repeated row-wise to match the dimensions of **M**. Principal components of **G** were computed using the *eigen* function in R (R Core Team, 2021).

To better understand the contribution of program differences to genetic variability, F_*ST*_ was estimated for each marker on a program and program group population basis. Following Cockerham (1973), estimation of F_*ST*_ was approached as the relative proportion of allelic variation associated with population labels relative to that associated with individual lines and genetic variability within lines. To accommodate different population sizes and allow for estimating population structure at both the group level (SunGrains^®^ versus non-SunGrains^®^) and program level, individual alleles *y*_*h*_ for each marker *h* were treated as a response in the nested model:

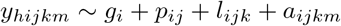

Where *y*_*hijkm*_ represents a numeric vector of allele calls for SNP marker *h*, two each from diploid individual *k* within breeding program *j* and breeding program group *i*. The proportion of variance associated with the within-line residual *a*_*ijkm*_ represents relative heterozygosity and can be used to construct an estimator of inbreeding 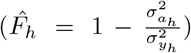. In this case however, lines are derived from mostly-inbred individuals with some residual heterozygosity, and allele calls produced by GBS are estimates of the underlying true genotype. As a result, 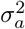 represents some combination of heterozygosity, within-line genetic heterogeneity, and genotyping error. Proportion of variance associated with the line variance *l*_*ijk*_ is taken to represent the relative contribution of between-line differences within programs to allelic diversity, while the proportion of variance associated with breeding programs *p*_*ij*_ and groups of breeding programs *g*_*i*_ indicate the relative importance of each level of population structure. Genome-wide F_*ST*_ associated with the group level is then taken as 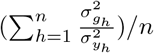 across *n* markers, while that associated with the program level is taken as 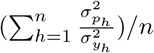. REML estimates of variance components were calculated for each marker using Asreml-R (Butler et al., 2017). Despite filtering markers on an overall heterozygosity of 0.10, for rarer alleles error-prone GBS markers might still overestimate the proportion of lines heterogenous for the allele and inflate the proportion of overall allelic variance associated with within-line variation. To remove these markers prior to calculation of F_*ST*_ statistics, the proportion of inbred lines expected to be heterogenous for a given marker was calculated based on the assumption that non-doubled haploid (DH) lines were derived at the F_4_ generation, such that the probability of a heterozygous marker call after the F_4_ generation 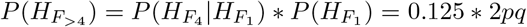. Markers were removed if the ratio between observed and expected heterozygosity in non-DH lines was greater than two (Supplemental Figure 1, Supplemental Figure 2).

### Environmental data

Weather data for the GPS coordinates associated with each site in each year were downloaded from NASA POWER using the nasaPower R package (Sparks, 2018). Within each site-year, principal grain fill periods were defined for each observed heading date by taking the time from the day after heading through 22 days. Additional variables were collected from twenty to thirty days after heading to test optimum grain filling periods for each weather variable. Literature suggests an effect of awns in improving photosynthesis during grain fill in drought and heat stress conditions as a result of better heat conductance, increasing water use efficiency with less transpiration, and later senescence. Therefore, data on max temperature, precipitation, wind speed, and relative humidity were collected. Collected weather variables across time points were averaged within each grain fill period associated with a heading date in a site-year.

### Testing and predicting weather effects

To estimate effects of *B1* on grain yield, a mixed linear model was fit to allow for the estimate of interactions between *B1* and environmental variables while controlling for effects of population structure and its interaction with environmental variables. Weather variables were calculated relative to heading date to allow for a range of variables within each site-year associated with each genotype’s mean heading date. Mean yield values for each line *i* were computed within each trial *j* and site-year *k* as *y*_*ijk*_ by collaborators prior to data aggregation. For awn status *a* of line *i*, matrix of relationship matrix principal components **Q** composed of *n* vectors *q* of components 1 through *n* with values for each line *i*, and matrix of weather variables **W** composed of *m* vectors *w* of different weather values for each line *i* within trial *j* in site-year *k*, the data was analyzed using the model:

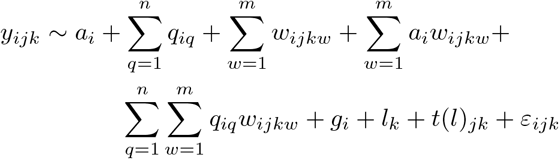

Where *g*_*i*_ represents the genotype effect of line *i* fit with the realized relationship matrix 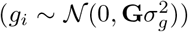, *l*_*k*_ represents the random effect of the *k*th site-year with 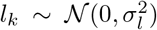, and *t*(*l*)_*jk*_ represented the nested random effect of the *j*th trial within the *k*th site-year with 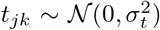. The residual *ε*_*ijk*_ is drawn from 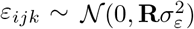, where the diagonal of **R** contains separate variances for different trials to accommodate heteroge-neous variances across trials 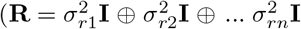 for *n* trials). As the model is fit in two stages, assumption of heterogeneous residuals represents an assumption of trial-to-trial differences in genotype by environment interaction variances and error variation not accounted for by replication (that is, deviation of genotype effects within each environment from overall genotype effects, and deviations of our estimates of genotypes within environments from the ”true”values that would be recovered with infinite replication). Practically, assumptions of heterogeneous trial variances weighs data from higher-variability trials less when estimating parameters. The model was fit in Asreml-R, and significance of main fixed effects was computed via the Wald test (Butler et al., 2017).

Differential adaption of lines from different breeding programs to different climates may relate to genetic divergence between programs as captured in measures of population structure, but may also relate to differences in selection environments between lines (Boyles et al., 2019). In the simplest case, we might expect a line submitted by the University of Georgia breeding program to be better adapted to heat stress than a full-sib line from the same cross submitted by the University of Maryland breeding program due to within-family selection in more heat-stressed Georgia environments, even if both lines would have similar principal component scores. Breeding programs in more southern environments have tended to prefer selecting awned lines, and northern programs awnless lines. This preference is due in part to the perceived better adaptation of awned lines to southern environments. Disentangling the effect of awns is complicated by the fact that lines from programs with overall better heat tolerance are more likely to be awned, while some component of that tolerance may itself relate to the greater frequency of awns in those programs. As a check, a separate, more conservative model was fit with a single detected weather variable *w* to assess the significance of interactions detected in the initial model after accounting for programs’ differential adaptation to those same stresses. Instead of matrix of principal components **Q**, a design matrix whose levels relate to individual breeding programs *P* was used to fit program-specific slopes on the detected weather variable *w* :

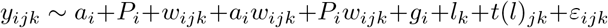

Where as before genotype effect of line *i g*_*i*_ was fit with the realized relationship matrix 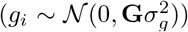, and residuals *ε*_*ijk*_ were drawn from 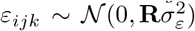. Unlike the initial model, the estimated slopes for individual programs on the weather variable *w* will include differences in programs’ response to that variable resulting from differences in the frequency of *B1*. As such, this model represents a lower bound on the magnitude of the reaction norm for *B1* by only using within-program differences in awn status in its estimation.

For both models, estimates of the fixed effects were used to derive reaction norms for allele effects of *B1* in interaction with significant environmental variables. Reaction norm equations were used to estimate allele effects of awns in each site-year, based on values for environmental variables observed in those site-years. Assuming that trial locations within states are representative of breeders’ target environments, site-year effects were grouped by state and averaged to estimate mean allele effects of awns within each state.

## Results and discussion

### Population structure of programs

To be useful in modeling the relative performance of lines from different programs under different environmental conditions, the principal components of the realized relationship matrix should capture genetic differences between programs resulting from patterns of parental selection. Visualization of primary principal components allows for the detection of severe population stratification, which would prevent accurate estimation of allele effects. The first four PCs of **G** together captured 14% of the total variation in relationships between 1,371 genotypes. Public programs were separated by these first four components (Fig. 1), but without severe stratification. This may be expected from the structure of wheat breeding programs in the Southeast US, which are distinct but cooperate and regularly share germplasm. Averaging the values of all individuals within each program shows that the first PC separated out the mid-Atlantic programs (MD, VA, KY, NC) from southern programs (FL, LA, TX, GA, SC), while the second PC separated programs within the mid-Atlantic and southern groups (Fig. 1). The fourth PC also effectively separated the most southern programs that may be more heat-stress resistant (FL, LA, and TX) from other germplasm. Discrepancies from previous studies using this germplasm (e.g. Sarinelli et al. (2019)), which showed the principal components being largely reflective of the presence or absence of SNP-dense translocations, result from LD thinning of the marker matrix performed prior to estimation of the realized relationships between lines.

**Figure 1:**
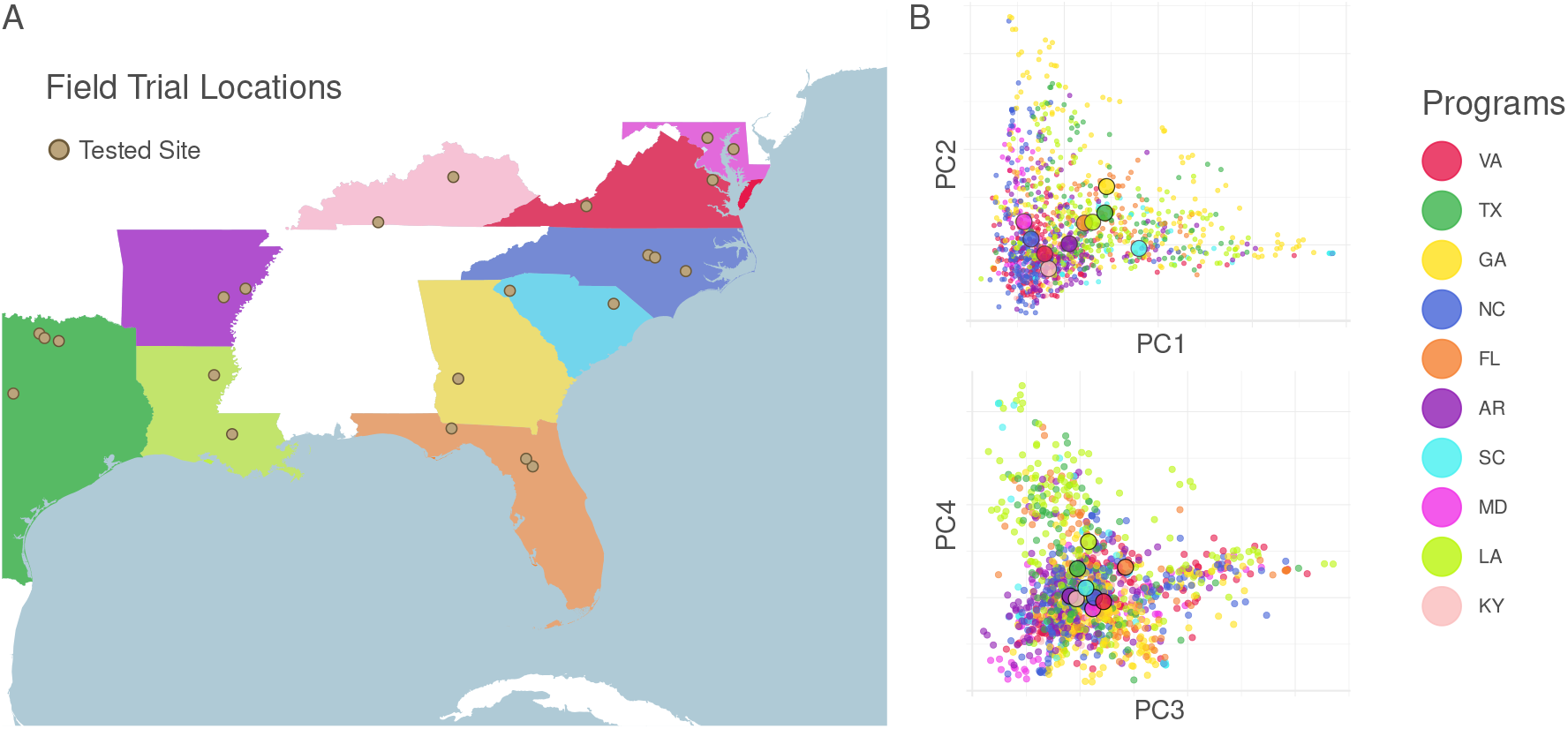
Geographic and genetic distance between public programs. Public-sector breeding programs contributing germplasm and phenotypes to tested nurseries are colored, and field site locations labeled (A). The distance between individuals in the space of the first four principal components of the realized relationship matrix **G** reflect geographic separation of the programs (B).

A measure of F_*ST*_ calculated by analysis of the components of allelic variance for each assessed SNP supported the partial population differentiation observed in the analysis of principal components (Table 2). Some differentiation was observed among breeding programs (0.016 for program group, 0.039 for program within group), although much less than the differentiation reported among market classes in USA wheat (Chao et al., 2007). A genome-wide scan reveals some genomic regions whose frequencies are differentiated by program structure (Supplemental Figure 3). Among these is *B1*, where population structure is substantial at both the group level (0.109) and program level (0.174). At the same time, the combined proportion of allelic variance associated with both levels of population structure is much less than that associated with individual lines (0.615), indicating that the majority of variation for the presence of *B1* is within populations.

**Table 2:** Components of allelic variance. The relative proportion of variance in allele calls 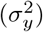 related to different levels of population structure (Group, member or non-member of SunGrains^®^; program, breeding program within group). Values are given for *B1* and for the mean of all remaining markers.

| Prop. $\sigma_y^2$ | *B1* | Genome-Wide |
| --- | --- | --- |
| Group | 0.109 | 0.018 |
| Program | 0.174 | 0.041 |
| Line | 0.615 | 0.812 |
| Within-Line | 0.102 | 0.130 |

### Environmental factors relevant to awn yield effects

Weather variables were selected based on their role in generating heat stress or altering transpiration rates: maximum temperature, precipitation, specific humidity, and wind speed information were obtained for the growing season of each site-year. An initial grain fill window of 22 days after heading was assumed for initial testing of the importance of variables, and the heading date of each plot in each site-year was used to average weather variables over that window. Of the tested variables, only maximum temperature was found to have a significant interaction with awn status after controlling for population structure. When testing possible alternative grain fill periods longer and shorter than 22 days, the grain fill period of 23 days produced the strongest interaction between average daily maximum temperature and *B1* 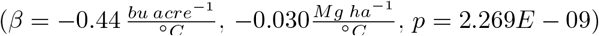. The first four principal components of **G** reflecting genetic distances between programs also interacted significantly with maximum temperature during grainfill (PC1, *p* = 1.61*E* − 03; PC2, *p* = 1.35*E* − 04; PC3, *p* = 1.98*E* − 07; PC4, *p* = 1.66*E* − 04). Re-testing the interaction with fixed effects for program origin instead of principal component scores also resulted in a highly significant, but smaller, interaction between temperature and *B1* (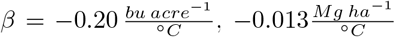, *p* = 1.767*E* −09 and between temperature and program of origin (*p* < 2.2*E* − 16), indicating a strong effect of awns in conferring heat tolerance beyond the different frequencies of *B1* in different programs. That the coefficient of the interaction effect is substantially smaller in the program model suggests that some component of programs’ differential adaption to temperature stress may include having a larger proportion of awned lines. At the same time, the larger coefficient estimate in the principal component model may result in part from covariance between *B1* frequency and program origin. The more conservative program model was chosen for subsequent analyses, with the understanding that the real effect of awns may be larger than estimated.

Both awned *b1* and awnless *B1* lines were lower yielding in environments with higher maximum temperatures during grain fill, but the decrease in yield was less severe for the awned lines than for awnless lines (Fig. 2). The lack of association between lower precipitation and improved yield for awned lines may suggest that direct cooling of the wheat head is the primary driver of improved yield for awned lines in heat-stressed conditions, but does not necessarily suggest that drought stress is unimportant physiologically for generating a yield effect of awns. Precipitation received during grain fill is only one of the drivers of drought stress experienced by plants during grain fill, as increasing temperatures may increase evapotranspiration and dry soil more rapidly. From the perspective of breeding for target environments, however, to what extent improved yield in heat-stress conditions for awned lines results from heat stress per se or drought stress driven by increasing temperatures is less important than knowing that awned lines out-perform awnless in those conditions. Increasing performance of *B1* awnless lines in cooler, higher-yielding environments may relate to the increase grain number associated with awn suppression. This increase may relate to minimizing photosynthate diversion to non-reproductive tissues at a critical developmental stage, or may have to do with other downstream effects of an increase in *B-A1* expression. While *B1* awn suppression may lower yields in poor environments, it may allow plants to take advantage of good growing conditions with less source limitations.

**Figure 2:**
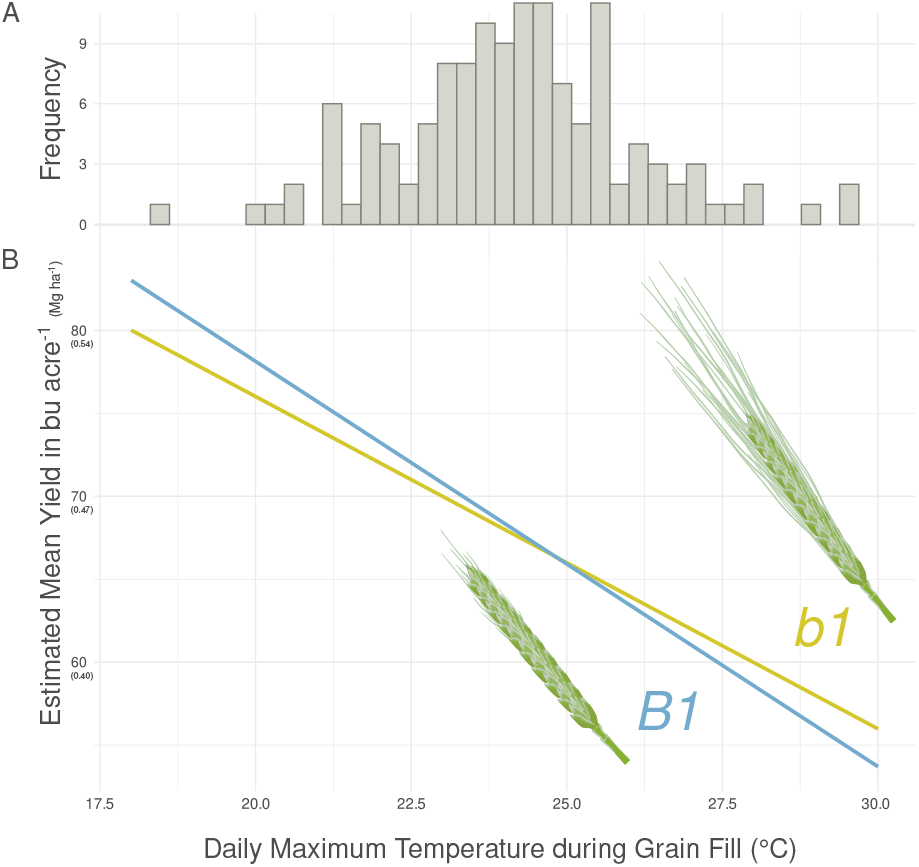
Interaction between *B1* and max grain fill temperature. The average maximum daily temperature during grain fill at each site-year was 24.1 °*C* (A), with a high maximum grain fill temperature of 29.8°*C* (Gainesville, FL 2017), and a low of 18.5°*C* (Warsaw, VA 2020). Estimated reaction norms for the conditional yield effect of the *B1* awn suppressor (B). *B1* awn suppression was favorable or neutral in most environment, but was associated with reduced yield in heat-stressed environments.

### The overall effect of awns on yield varies by environment, year, and germplasm

Historic data on testing environments were used to estimate the yield effect of awns in each site-year. Reaction norms for *B1* estimated in the conservative mixed model that accounts for program by temperature effects were combined with the heading-date based grain fill mean daily maximum temperatures to estimate environment-specific yield effects of *B1* (Fig. 3). While the average estimated effect of awns was slightly negative (−0.28 bu acre^−1^, -0.018 Mg ha^−1^), the effect within site-years varied with temperatures from a decrease of 2.8 bu acre^−1^ (−0.18 Mg ha^−1^, Warsaw, VA in 2020), to an increase of 2.1 bu acre^−1^ (0.14 Mg ha^−1^, Gainesville, FL 2017). Breeders generally select field sites representative of their target environments, but the proportions of warmer to cooler environments sampled by collaborators through the trials in this data set likely does not accurately represent the relative proportions of those environments in real wheat acreage. Nevertheless, to try to understand the within-state patterns of *b1* yield effects, effects on all field trials conducted within each state were averaged over years (Table 3). Overall positive effects of awns on yield were observed only in sites in Florida and Texas. At the same time, the majority of states with an overall negative yield effect of awns had years with a positive awns effect more than a quarter of the time. While the states with the overall largest advantage for *b1* and the largest advantage for *B1* were the most southern (FL) and most northern (MD) states respectively, awn effects did not scale exactly with the overall climate of states. Despite being the state with the overall coldest climate, Kentucky had the third-highest yield effect of *b1*, and an overall positive effect of *b1* in 40% of years (Table 3). This likely results from the observation that Kentucky mean heading dates were on average the latest of any state (128 days), pushing grain fill into late May when temperatures rise. Conversely, South Carolina had one of the largest overall negative yield effects of awns, potentially relating to its average early mean heading date (97.8). Because heading date is itself under the control of environmental conditions, the effect of awns on yield may not reflect overall climactic differences between regions. At the same time, the effect of awns on yield will vary within a population, reflecting differences in flowering habits of lines.

**Figure 3:**
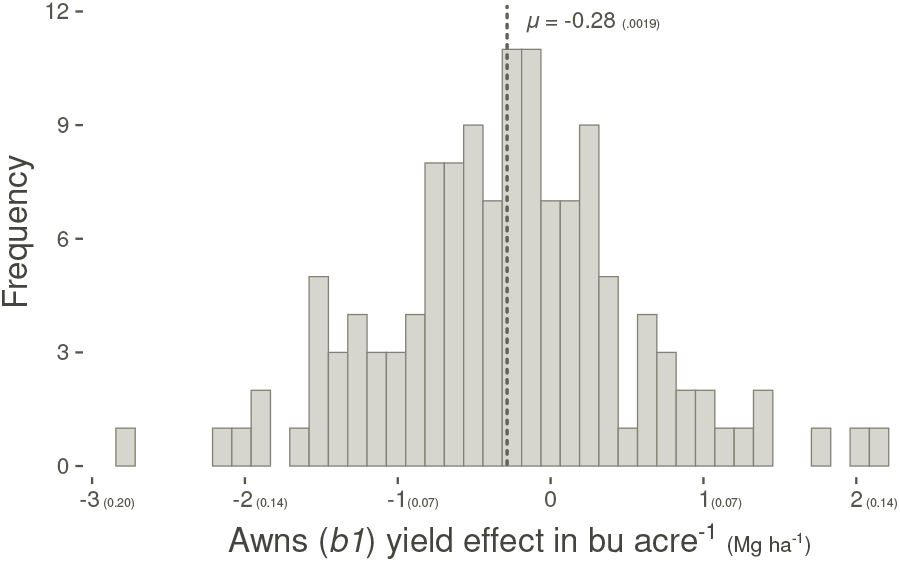
Distribution of estimated per-site *b1* yield effects. The mean estimated yield effect of awns is slightly negative in sampled environments, but environmental conditions associated strong positive and negative yield effects are also observed.

**Table 3:** Mean observations and estimated effects. States ranked by the mean effect of awns (*b1*) in bu acre^−1^ and Mg ha^−1^ (in parentheses) across all site-years. Mean values for heading date (HD), yield, and max grain fill temperature are given, along with the percentage of trials for which the estimated effect of awns was positive (% Trials +). The percentage of trials where the estimated effect of awns was positive can be substantial even in states where the mean effect was negative due to year-to-year variability in heading date and temperature.

| State | $\mu_{HD}$ | $\mu_{Yield}$ | $\mu_{GFMaxC}$ | $b1$ Effect | % Trials + |
| --- | --- | --- | --- | --- | --- |
| FL | 95.4 | 56.66 | 26.98 | 0.95 (0.064) | 77 |
| TX | 99.5 | 60.36 | 24.98 | 0.07 (0.004) | 60 |
| KY | 128.1 | 76.92 | 24.52 | -0.13 (0.009) | 40 |
| LA | 92.9 | 56.94 | 24.42 | -0.18 (0.012) | 20 |
| GA | 96.4 | 73.83 | 24.19 | -0.28 (0.018) | 25 |
| AR | 101.7 | 58.57 | 23.94 | -0.39 (0.026) | 08 |
| NC | 104.6 | 66.80 | 23.55 | -0.56 (0.038) | 33 |
| SC | 97.8 | 81.13 | 23.30 | -0.67 (0.045) | 33 |
| VA | 118.8 | 77.05 | 23.24 | -0.70 (0.047) | 30 |
| MD | 125.9 | 81.04 | 22.66 | -0.95 (0.064) | 00 |

Breeders may consider the effect of awns on both overall yield and on year-to-year variability in yield. While in a majority of sampled environments the *b1* allele for awn presence was associated with negative yield, yields in these environments were overall higher than environments where *b1* had a positive effect. If awnless lines have greater yield potential in favorable conditions, but suffer in stressed environments, then their overall variability will be higher. In environments where the yield effect of awns is sometimes negative and sometimes positive, growers may prefer awned lines with a higher yield ”floor”, even if they have lower overall yield potential.

Predicted variability in the yield of *B1* versus *b1* lines within each state was estimated using the site-year estimates of awn effects based on weather data and heading dates (Table 4). While Virginia was the state with the second highest yield advantage for awnless lines, high variability in grain fill temperatures resulted in a much higher estimated yield variability for awned lines compared to awnless. This contrasts with Arkansas, where *B1* awnless lines were predicted to be overall higher-yielding but only marginally more variable than awned lines. Given that growers have little power to predict the environmental conditions of next year’s growing season, selecting for traits like awns that help produce consistent yields under a range of conditions may be valuable in environmentally variable locations.

**Table 4:** Site-year variance in observations and estimated *B1* effects. Means for between-site-year observed variances of heading date (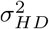, Julian days) and grain fill temperature (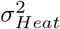, °*C*) are compared to yield variance in estimated means of lines with awns (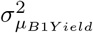, bu acre^−1^, Mg ha^−1^ in parentheses) and without awns (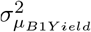, bu acre^−1^, Mg ha^−1^ in parentheses) States are ranked according to the difference in variances of estimated *B1* and *b1* yields (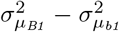, bu acre^−1^, Mg ha^−1^ in parentheses), a measure of the estimated benefit of awn lines in reducing year-to-year yield variance.

| State | $\sigma_{HD}^2$ | $\sigma_{Heat}^2$ | $\sigma_{\mu_{B1Yield}}^2$ | $\sigma_{\mu_{b1Yield}}^2$ | $\sigma_{\mu_{B1}}^2 - \sigma_{\mu_{b1}}^2$ |
| --- | --- | --- | --- | --- | --- |
| VA | 77.5 | 5.06 | 30.3 (0.137) | 20.3 (0.092) | 10.0 (0.045) |
| FL | 60.3 | 3.32 | 19.8 (0.090) | 13.3 (0.060) | 6.5 (0.030) |
| NC | 57.7 | 2.89 | 17.3 (0.078) | 11.6 (0.052) | 5.7 (0.026) |
| MD | 130.7 | 2.92 | 17.4 (0.079) | 11.7 (0.053) | 5.7 (0.026) |
| TX | 147.2 | 2.74 | 16.4 (0.074) | 11.0 (0.050) | 5.4 (0.024) |
| SC | 72.5 | 2.53 | 15.1 (0.068) | 10.2 (0.046) | 4.9 (0.022) |
| KY | 17.2 | 1.64 | 9.8 (0.044) | 6.6 (0.030) | 3.2 (0.014) |
| LA | 50.9 | 1.58 | 9.5 (0.043) | 6.4 (0.029) | 3.1 (0.014) |
| GA | 36.0 | 0.96 | 5.7 (0.026) | 3.8 (0.017) | 1.9 (0.009) |
| AR | 147.6 | 0.73 | 4.4 (0.020) | 2.9 (0.013) | 1.5 (0.007) |

## Conclusions

The obvious visual difference between awned and awnless lines has made *B1* one of the most studied genes in crop genetics. Making sense of conflicting results on the importance of this gene in contributing to yield variation, potentially in an environment-specific manner, has been limited by geneticts’ ability to sample a sufficient number of diverse environments and genotypes to make real inferences on its yield effect. Here, we present the largest study yet on this gene, and find that its effect on yield varies in relation to heat stress during grain fill. We estimate that while awns have a slight negative effect across years for all states in our data set except Texas and Florida, on a year-to-year basis the effect of awns may be positive or negative, depending on heat stress during grain fill. Overall estimated effects suggest that *B1* awn suppression provides a slight yield improvement in high-yielding environments, but will lower yields relative to awned lines in lower-yielding, heat stressed environments. Breeders already account for these effects and select for awned lines; for this reason, the conservative model used in these predictions, which estimated individual reaction norms for each program of origin, may under-estimate the overall effect size of *B1* and its interaction with heat stress. Considerations on selecting for or against *B1* relate not just to its yield effects in controlled environments; awns protect against herbivory, limiting crop damage from deer, and growers may prefer awned varieties for aesthetic reasons or due to the historic success of certain awned varieties. Despite this, breeders targeting environments that rarely experience grain fill heat stress may consider selecting awnless *B1* lines due to the better performance of awnless lines in those conditions. Breeders targeting environments where awns have a small average negative yield effect may still consider selecting awned lines for their higher yield ”floor”and lower year-to-year yield variability. As both overall temperature and year-to-year variability in heat stress increase, breeders in these marginal environments may also consider preferentially selecting awned lines for future climates.

In our germplasm, *B1* interacts with heat stress to generate differential effects on yield in different environments. At the same time, the effect does not scale linearly with the overall climate of testing locations, due to differences in plant phenology – *B1* will have a different effect in plants that head early and escape heat stress than those that flower later. *B1* was chosen because of its ease to characterize and history as a major gene implicated in resistance to heat stress, but any genes that confer abiotic stress tolerance without altering phenology will likely behave similarly. Understanding the genetic basis of plant adaptation to changing climates may necessitate first modeling the genetic basis of and relationship between plant development and stress avoidance.

## Supplementary data

Supplementary file 1 contains additional figures relating to analysis of marker heterozygosity and *F*_*st*_ calculations.

## Acknowledgements

The authors thank the staff of the USDA-ARS Plant Sciences unit for help in collection of field and sample preparation of genotype data. Thanks to Zachary Winn and Nicole Choquette for feedback on methods used in the manuscript. The authors also thank staff of cooperating breeding programs for planting and phenotyping of cooperative yield trials. Weather data was obtained from the NASA Langley Research Center (LaRC) POWER Project funded through the NASA Earth Science/Applied Science Program.

## Author’s contributions

ND prepared genotype data, performed all analyses, and wrote the initial draft of the manuscript. ND, MG, JPM, JBH, and GBG conceived of the analyses. ND, MG, JPM, JBH, and GBG edited the manuscript. JL and BPW organized and prepared the phenotypic data. JPM, REB, MM, MAB, ES, RS, AI, DAVS, VT, NS, and SAH planted and managed yield trials, collected phenotypes, and performed initial within-site analyses. KH and JS designed and optimized the genotyping pipeline and prepared all samples for sequencing.

## Data availability

Breeding program data, including phenotypes and genotypes associated with individual programs’ lines, may be available on a program-by-program basis by request.

