## Supplemental data 1 for "Bearded or smooth? Awns improve yield when wheat experiences heat stress during grain fill"

**Supplemental Figures**

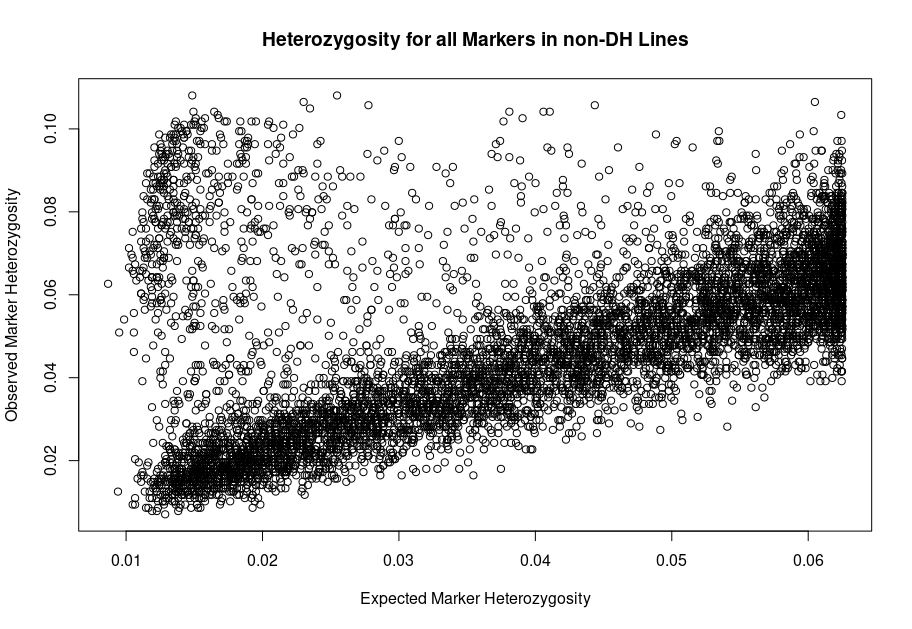
**Supplemental figure 1 – Expected versus observed marker heterozygosity.** Observed marker heterozygosity in inbred lines is expected to be a combination of genetic heterogeneity due to heterozygosity at the F generation from which lines were derived, genotyping error, and some potential residual heterozygosity. Expected marker heterozygosity is calculated assuming lines are derived at the F4 generation as the product of a chance a marker is heterozygous in the F4 given that the marker was heterozygous in the F1 (0.125) times the chance of an F1 being heterozygous (assumed to be HW value for heterozygotes at 2pq, or 2 * MAF * (1-MAF). Expected marker heterozygosity tends to be fairly predictive of observed heterozygosity, except for some outliers that were excluded in subsequent analyses.

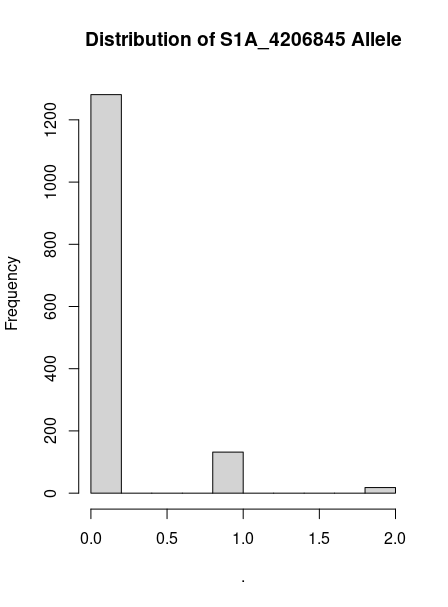

**Supplemental figure 2 – Example of outlier allele.** Outlier heterozygous allies persist despite filtering on minor allele frequency (MAF < 0.05) and heterozygosity (HZ < 0.10) when rarer alleles above MAF of 0.05 are persistently miscalled as heterozygotes.

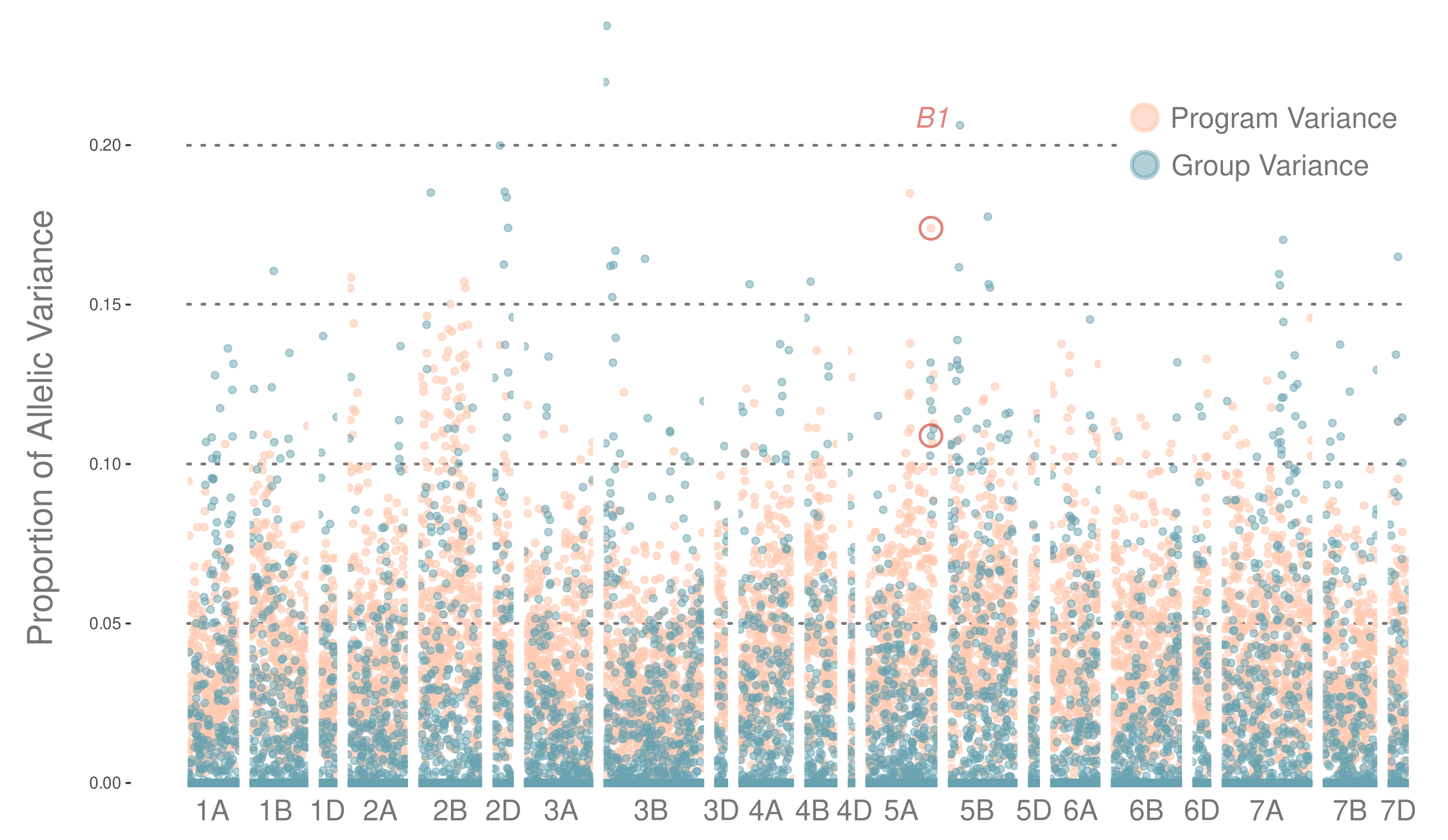
**Supplemental figure 3 – Genome-wide FST.** Proportion of variance associated with program-level population structure and group level population structure (SunGrains vs non-SunGrains) plotted for each marker. SNP marker S5A_698528417 linked to *B1* shows evidence of differentiation due to population structure, but to a similar degree as other chromosomal regions.
